# Extrachromosomal DNA is associated with decreased immune cell infiltration and antigen presentation, represents a potential cancer immune evasion mechanism

**DOI:** 10.1101/2022.02.04.479205

**Authors:** Tao Wu, Chenxu Wu, Xiangyu Zhao, Guangshuai Wang, Wei Ning, Ziyu Tao, Fuxiang Chen, Xue-Song Liu

**Author notes:** Correspondence: Xue-Song Liu, School of Life Science and Technology, ShanghaiTech University, 230 Haike Road, Shanghai 201210, China. These authors contributed equally to this work (Tao Wu, Chenxu Wu).

## Abstract

Extrachromosomal DNA (ecDNA) is a type of circular and tumor specific genetic element. EcDNA has been reported to display open chromatin structure, facilitate oncogene amplification and genetic material unequal segregation, and is associated with poor cancer patients’ prognosis. The ability of immune evasion is a typical feature for cancer progression, however the tumor intrinsic factors that determine immune evasion remain poorly understood. Here we show that the presence of ecDNA is associated with markers of tumor immune evasion, and obtaining ecDNA could be one of the mechanisms employed by tumor cells to escape immune surveillance. Tumors with ecDNA usually have comparable TMB and neoantigen load, however they have lower immune cell infiltration and lower cytotoxic T cell activity. The microenvironment of tumors with ecDNA shows increased immune desert, decreased immune enriched fibrotic types. Both MHC class I and class II antigen presentation genes’ expression are decreased in tumors with ecDNA, and this could be the underlying mechanism for ecDNA associated immune evasion. This study provides evidence that the presence of ecDNA is an immune escape mechanism for cancer cells.

## Introduction

The immune system plays a crucial role in the protection and fight against cancer cells^1,2^. Immunoediting, which includes three temporally distinct stages, termed elimination, equilibrium, and escape, has been proposed to explain the interactions between cancer cells and the immune system during the evolution of cancer^3–5^. The mechanisms responsible for the escape of tumor cells from immunosurveillance are not fully understood. Potential tumor intrinsic immune evasion mechanisms include: impaired antigen presentation machinery (such as B2M mutation, decreased antigen presentation gene expression^6–8^), overexpressed immune checkpoints or their ligands such as programmed death-ligand 1 (PD-L1) on cancer cells^9^. In addition, secreting of immune inhibitory cytokines, such as TGF-β, remarkably reshape the tumor immune microenvironment^10,11^.

Extrachromosomal DNA (ecDNA) is a type of tumor specific DNA element that is circular and about 1–3 Mb in size. Since the 1960s, double minute chromosomes have been observed in the metaphase spreads of human cancer cells^12^. Later these DNA elements without centrioles and telomeres are found to be circular, a few Mb in size, and their size but not their number is stable during the proliferation of cancer cells^13^. With the recent advance of sequencing and bioinformatics techniques, ecDNA has been found to be prevalent in various types of cancers, however ecDNA is rarely detected in normal tissues, suggesting the presence of ecDNA is a specific feature for some cancer cells^14^. EcDNA promotes accessible chromatin (open chromatin) formation, facilitates oncogene amplification, drives genetic heterogeneity, and is associated with poor prognosis in multiple types of cancer^15–17^.

Somatic DNA alterations are major determinants of cancer phenotypes, including immune phenotypes. EcDNA formation is a type of somatic DNA alteration. We hypothesize that ecDNA formation could be one mechanism for cancer cells to evade immune surveillance.

## Results

### EcDNA and tumor immune cell infiltration status

For this study, we select cancer patient samples with both WGS and gene expression data for analysis. The status of ecDNA in specific samples was determined based on WGS data as previously described^17^. In total, 1684 samples with ecDNA status and gene expression information are available for analysis (Supplementary Fig. S1).

First we investigate the correlation between the presence of ecDNA and tumor immune infiltration status. The immune infiltration status was determined using gene mRNA expression data. Multiple methods have been applied in the quantification of tumor immune status, including TIMER, CIBERSORT, Xcell, MCPcounter, Quantiseq and Estimate^18–23^. With different methods, tumors with ecDNA consistently show significantly decreased immune scores (Fig. 1a-c and Supplementary Fig. S2). Importantly, the cytotoxic T cell (CD8^+^) levels and cytotoxic scores are significantly decreased in tumors with ecDNA (Fig. 1d and Supplementary Fig. S3). The composition of different immune cells was calculated using gene expression data with multiple different methods, including marker gene-based methods (Xcell and MCPcounter) or deconvolution-based methods (Cibersort, Timer, and Quantiseq). Multiple types of immune cells including B cell, NK cell and T cell show significantly decreased composition in tumors with ecDNA in TCGA pan-cancer dataset as a whole (Fig. 2a), or in separate cancer types, such as STAD (Stomach adenocarcinoma), SKCM (Skin cutaneous melanoma), HNSC (Head and neck squamous cell carcinoma) (Fig. 2b and Supplementary Fig. S3).

**Figure 1.**
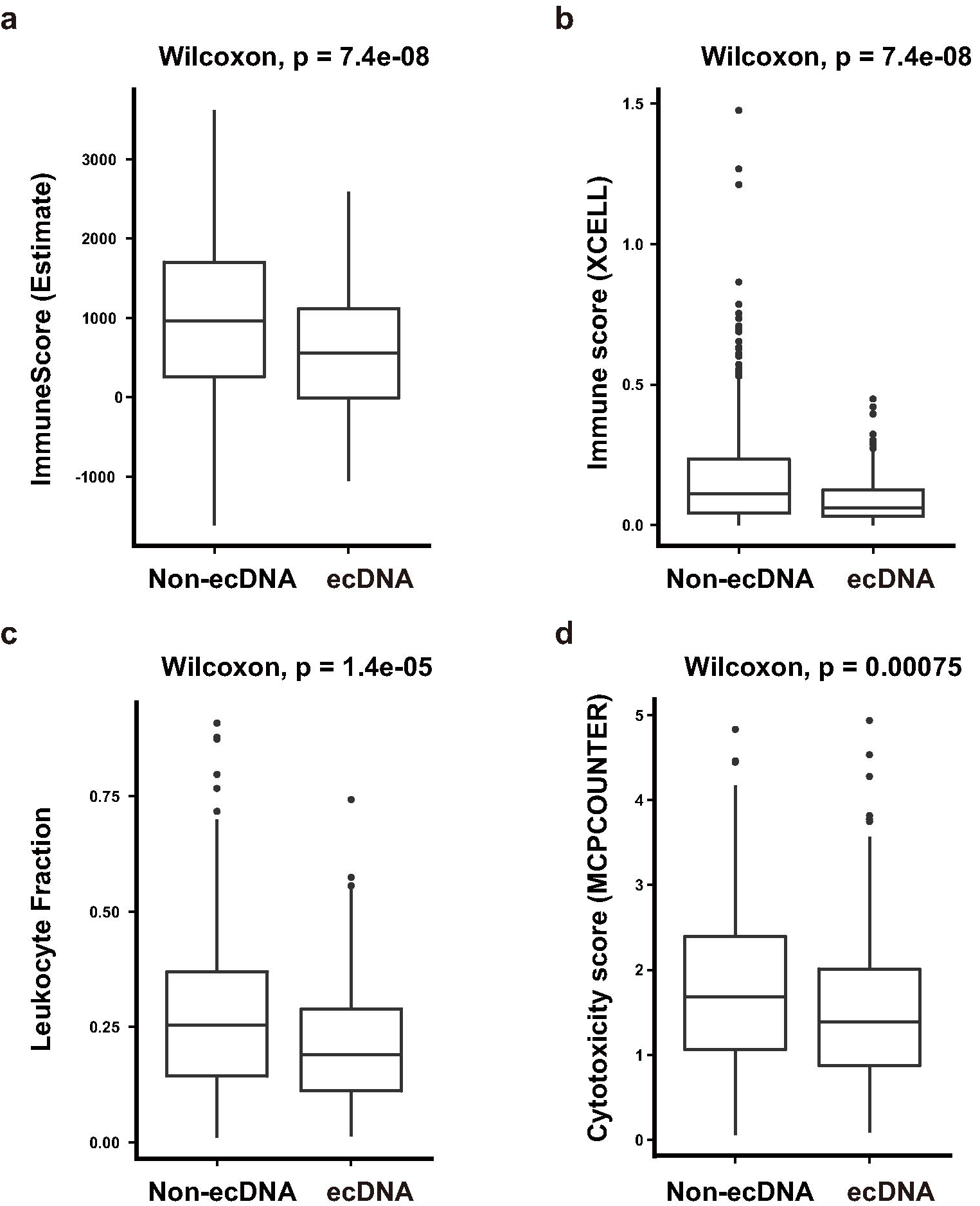
ecDNA and tumor immune infiltration scores. **a-d** Comparisons of immune infiltration scores quantified by different methods between tumors with and without ecDNA. **a** Estimate ImmuneScore; **b** XCELL Immune score; **c** Leukocyte fraction, **d** MCPCounter cytotoxicity score. Wilcoxon test p values are shown.

**Figure 2.**
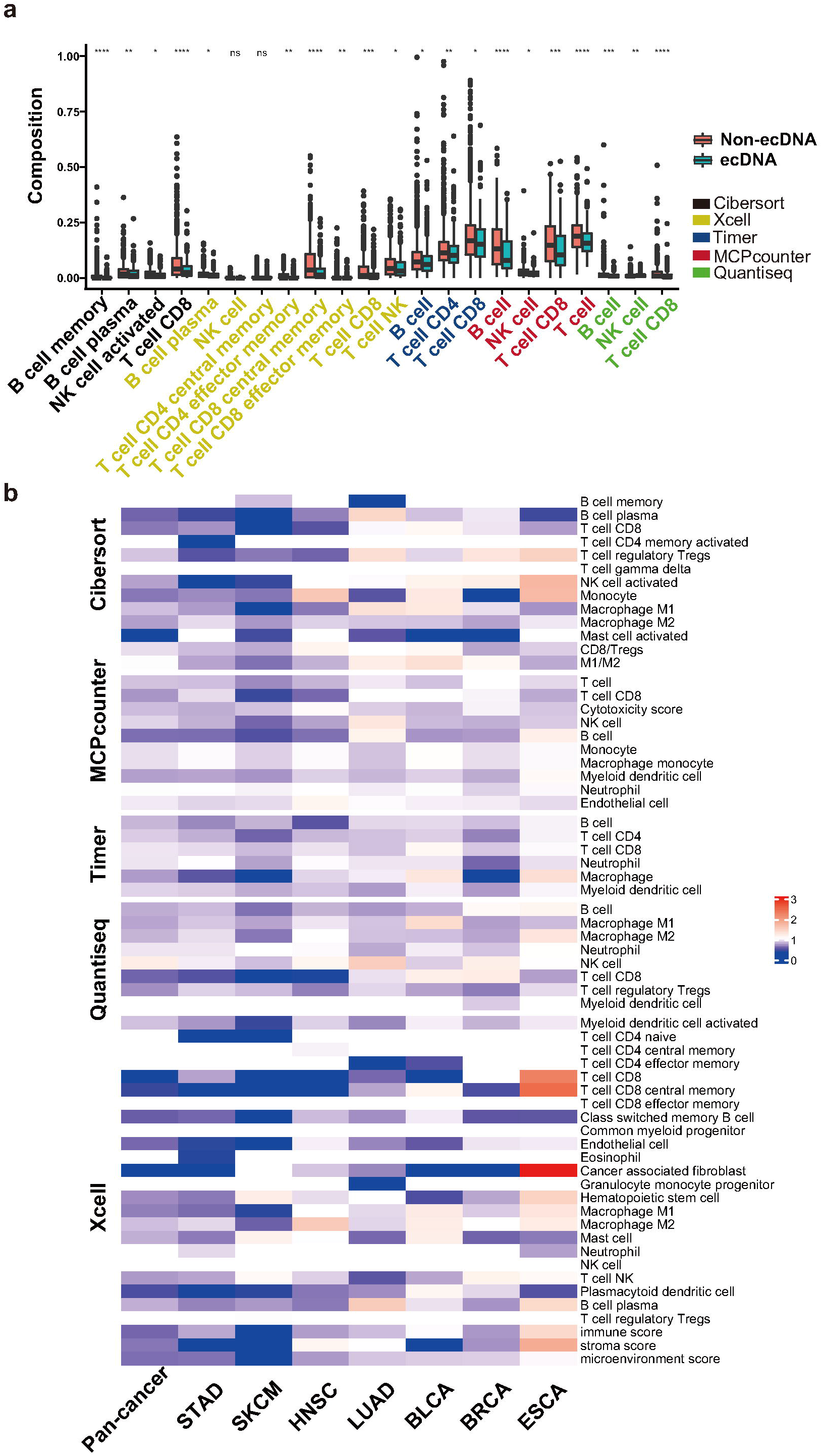
ecDNA and the infiltration of different types of immune cells. **a** Comparisons of the compositions of different types of immune cells between tumors with ecDNA and without ecDNA. The immune cell compositions have been quantified by five different methods, including Cibersort, Xcell, Timer, MCPcounter and Quantiseq. Wilcoxon test p values are shown. ns: *P*>0.05, *: *P*≤0.05, **: *P*≤0.01, ***: *P*≤0.001, ****: *P*≤0.0001. **b** Comparison of immune cell infiltration levels quantified by five different methods between ecDNA and non-ecDNA samples in different cancer types. Heatmap color indicates ratio of the median infiltration level for specific immune cell and specific cancer type between ecDNA and non-ecDNA samples. TCGA cancer type acronyms: STAD (stomach adenocarcinoma), SKCM (skin cutaneous melanoma), HNSC (head and neck squamous cell carcinoma), LUAD (lung adenocarcinoma), BLCA (bladder urothelial carcinoma), BRCA (breast invasive carcinoma), ESCA (esophageal carcinoma).

### EcDNA and tumor immune typing

Tumor immune typing was performed according to two known studies^24,25^. Thorsson et al used consensus clustering based on scored immune expression signatures to cluster cancer samples into six immune subtypes—wound healing, IFN-γ dominant, inflammatory, lymphocyte depleted, immunologically quiet, and TGF-β dominant^25^. In tumors with ecDNA, lymphocyte depleted type is up-regulated, while inflammatory and TGF-β dominant types are down-regulated (Fig. 3a). Bagaev et al used unsupervised dense Louvain clustering based on ssGSEA scores of 29 Fges (functional gene expression signatures) of immune and stromal related genes to cluster cancer samples into four distinct microenvironments: (1) immune-enriched, fibrotic (IE/F); (2) immune-enriched, non-fibrotic (IE); (3) fibrotic (F); and (4) immune-depleted (D)^24^. In tumors with ecDNA, fibrotic immune-enriched type of TME (IE/F) is dramatically decreased, while immune desert type TME (D) is significantly up-regulated (Fig. 3b).

**Figure 3.**
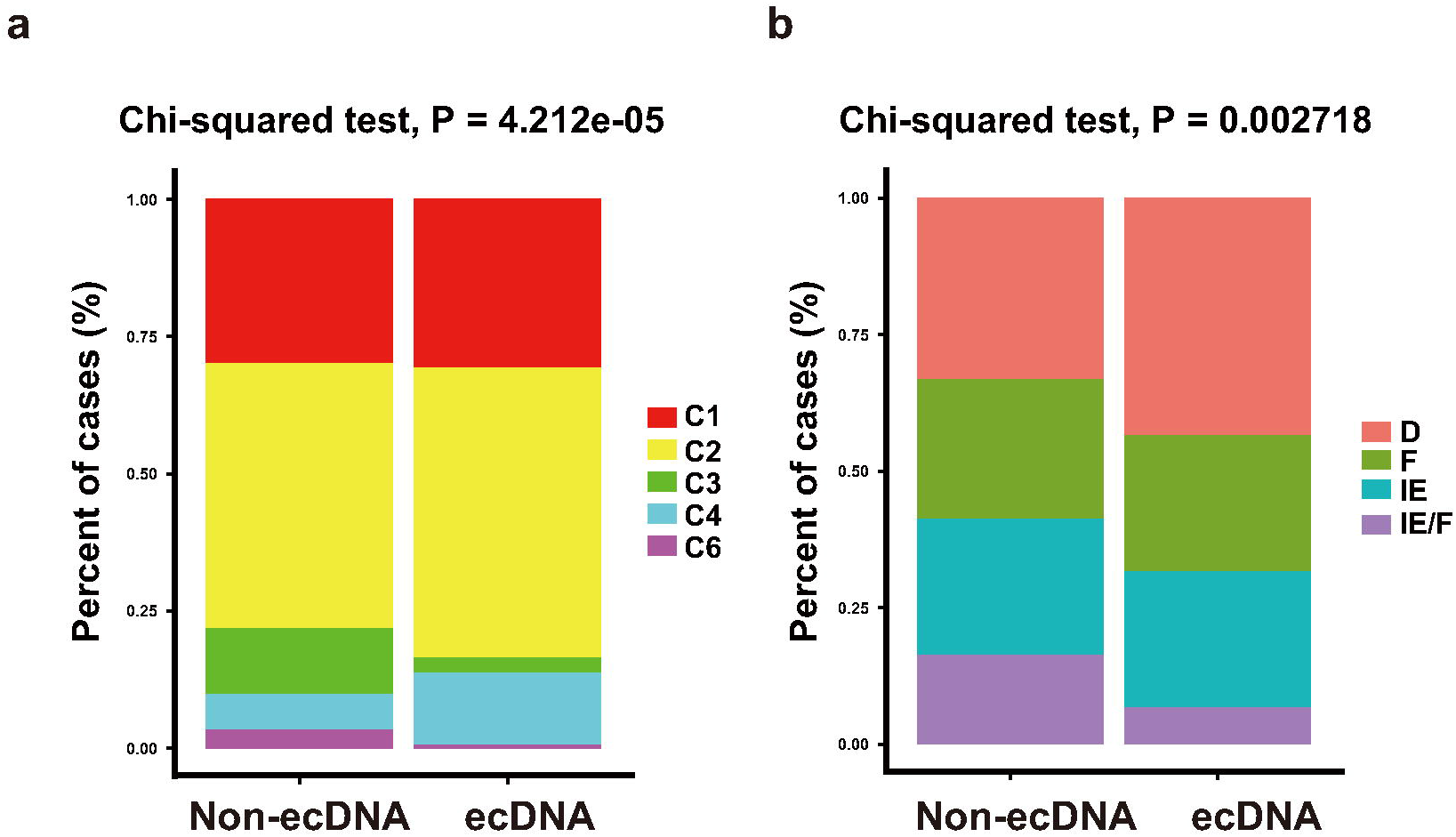
ecDNA and tumor immune typing. **a** TME classification in tumors with and without ecDNA according to Thorsson et al method. Chi-squared test p value is shown. C1: wound healing; C2: IFN-γ dominant; C3: inflammatory; C4: lymphocyte depleted; C5: immunologically quiet; C6: TGF-β dominant. **b** Immune type classification in tumors with ecDNA and without ecDNA according to Bagaev et al method. Chi-squared test p value is shown. D: immune-depleted; F: fibrotic; IE: immune-enriched, non-fibrotic; IE/F: immune-enriched, fibrotic.

### EcDNA and tumor immune escape

Expression of immune inhibitory immune checkpoint genes, such as PD-L1, CTLA4 is significantly down-regulated in tumors with ecDNA (Fig. 4a and Supplementary Fig. S4), suggesting the immune evasion of tumors with ecDNA is not through stimulating immune checkpoint signaling. This also implicates that immune checkpoint inhibitor therapy alone may not work in tumors with ecDNA.

**Figure 4.**
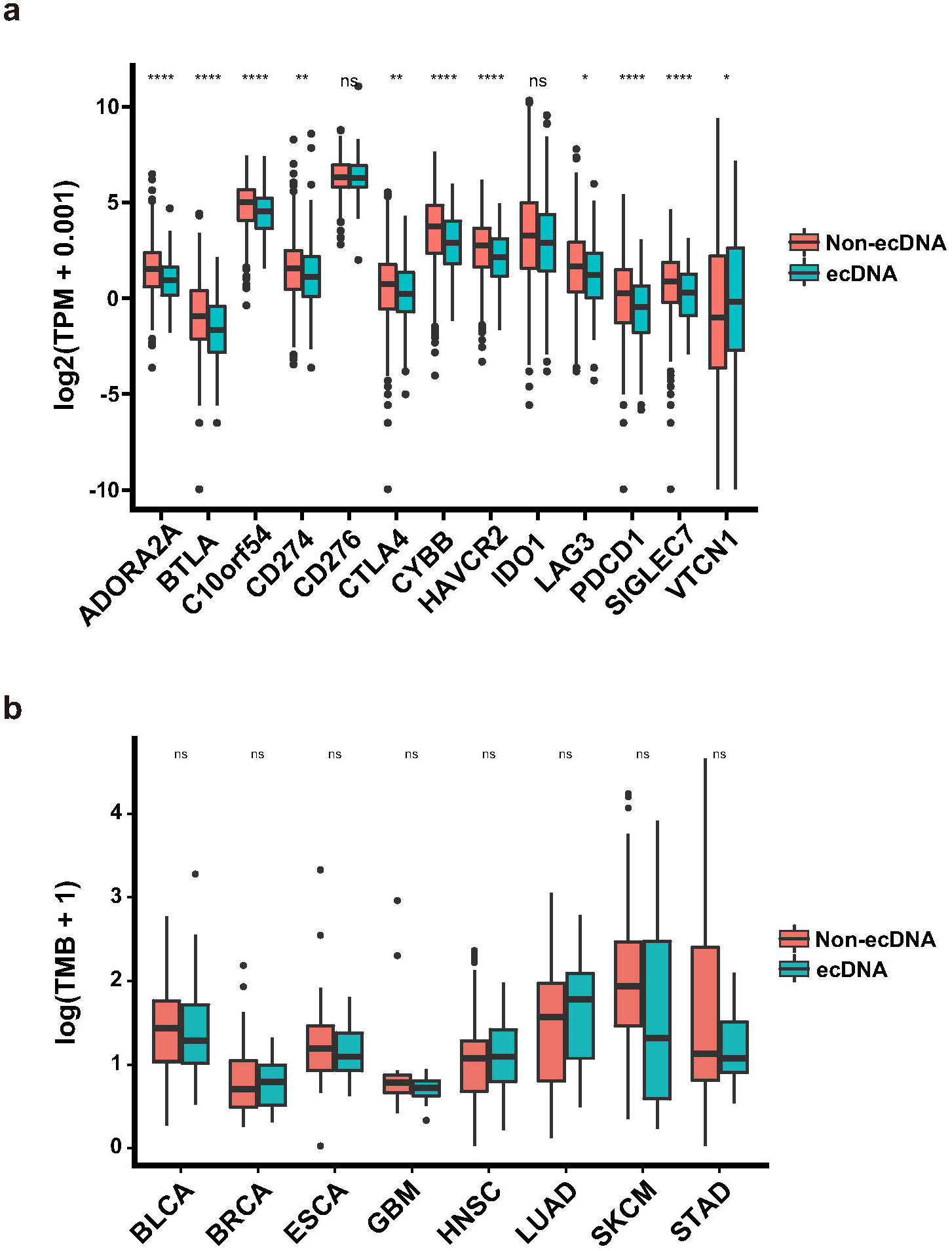
ecDNA and expression of inhibitory immune checkpoint genes and TMB. **a** Expression of inhibitory immune checkpoint genes in tumors with ecDNA and without ecDNA. Wilcoxon test *P* values are shown. **b** Tumor mutation burden (TMB) difference in different types of tumors with and without ecDNA. Wilcoxon test p values are shown. ns: *P*>0.05, *: *P*≤0.05, **: *P*≤0.01, ***: *P*≤0.001, ****: *P*≤0.0001. TCGA cancer type acronyms: STAD (stomach adenocarcinoma), SKCM (skin cutaneous melanoma), HNSC (head and neck squamous cell carcinoma), LUAD (lung adenocarcinoma), BLCA (bladder urothelial carcinoma), BRCA (breast invasive carcinoma), ESCA (esophageal carcinoma), GBM (glioblastoma multiforme).

### Antigen presentation and ecDNA mediated immune escape

Tumors with ecDNA show decreased immune cell infiltration, suggesting a decrease of immunogenicity in ecDNA-containing tumor cells. The immunogenicity of tumor cells determines the tumor associated immune response, and the antigenicity encoded by neoantigenic mutations is an important determinant of tumor immunogenicity^26^. Tumors with ecDNA show comparable TMB and neoantigen counts, suggesting a comparable antigenicity (Fig. 4b and Supplementary Fig. S5). This implies that the decreased immunogenicity of ecDNA-containing tumors was not caused by impaired antigenicity.

Antigen presentation efficiency is another important determinant of tumor immunogenicity^26^. The function of MHC class I antigen presentation pathway is to display peptide fragments of proteins from within the cell to cytotoxic T cells; MHC Class II molecules are normally found only on professional antigen-presenting cells such as dendritic cells, mononuclear phagocytes, some endothelial cells, thymic epithelial cells, and B cells. The antigens presented by class II peptides are derived from extracellular proteins. Expression of antigen presentation related genes, including MHC I, MHC II related genes, are compared between tumors with and without ecDNA (Fig. 5a and Supplementary Fig. S6). In tumors with ecDNA significantly decreased expression of MHC class I and class II genes are observed (Fig. 5a). Gene set enrichment analysis indicates MHC class I and class II related genes are significantly down-regulated in several cancer types (Fig. 5b). The impaired expression of MHC I and II related antigen presentation genes could be the mechanism underlying decreased immune infiltration in tumors with ecDNA.

**Figure 5.**
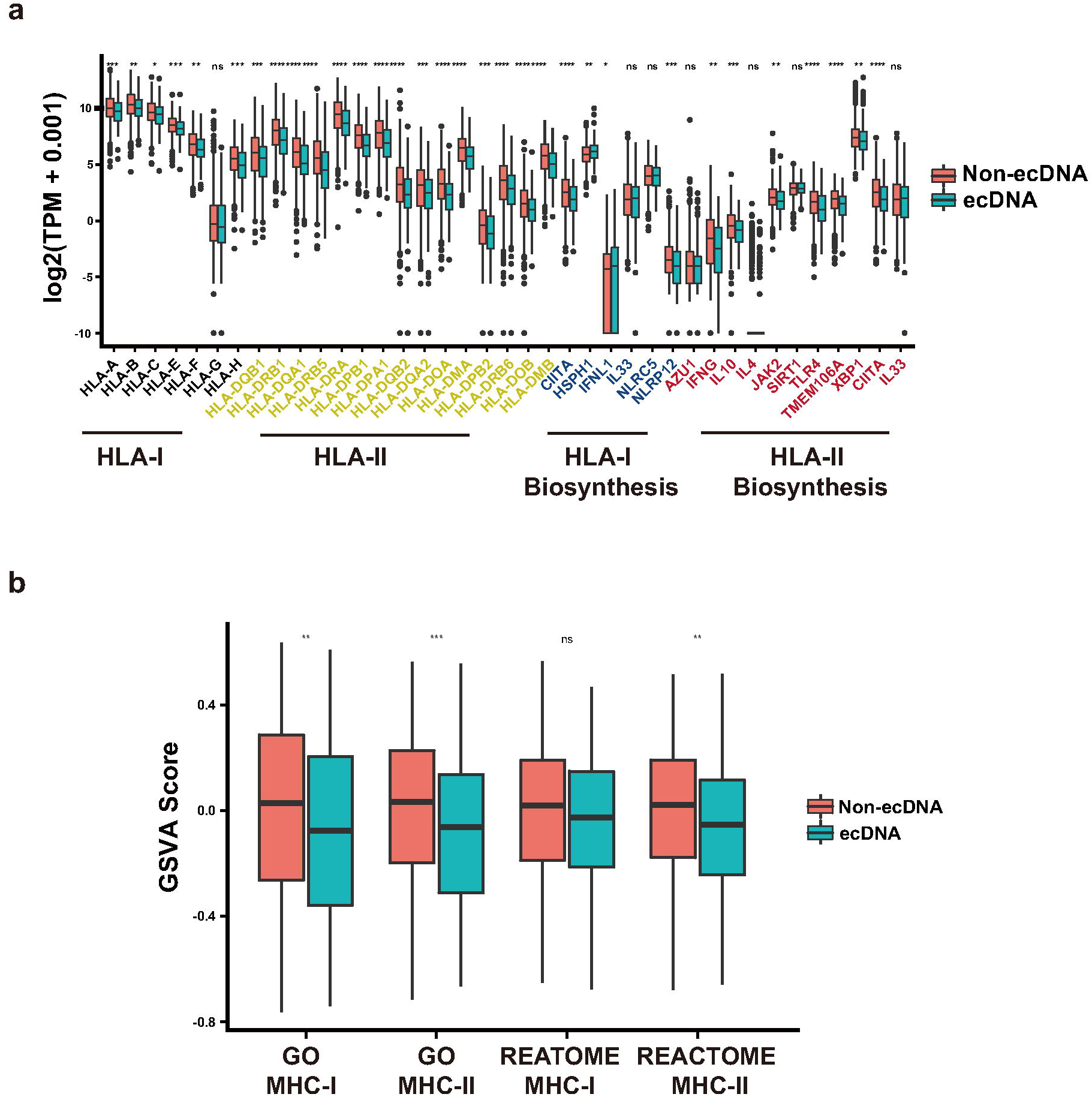
ecDNA and antigen presentation genes’ expression. **a** mRNA expression of MHC class I and class II antigen presentation related genes in tumors with and without ecDNA. Wilcoxon test p values are shown. **b** GSVA scores of MHC class I or class II antigen presentation genes in tumors with and without ecDNA. Wilcoxon test *P* values are shown. ns: *P*>0.05, *: *P*≤0.05, **: *P*≤0.01, ***: *P*≤0.001, ****: *P*≤0.0001.

## Discussion

Here we provide evidence to show that the presence of ecDNA is associated with decreased immune cell infiltration, decreased cytotoxic T cell percentage/composition, decreased expression of both class I and class II antigen presentation machinery genes. This analysis indicates that the presence of ecDNA could be one of the mechanisms employed by tumor cells to evade immune surveillance. EcDNA is preferentially detected in tumors, and less frequently in cultured tumor cells^27^. The immune selection pressure in tumors could be the underlying mechanism for this observation.

This study is based on gene expression data derived from bulk tumor samples, currently it is unclear if the gene expression differences happens in tumor cells or in the microenvironment immune cells or stromal cells. Consistently down-regulated antigen presentation related genes are observed in various types of tumors with ecDNA, and the functional consequence of these gene expression down-regulation in antigen presentation process need to be examined using experimental assays.

Based on this study ecDNA could directly induce tumor immune escape through down-regulating the expression of antigen presentation genes. Currently there are no experimental evidences supporting the alternative possibility that immunosuppressive microenvironment directly induces ecDNA formation. Potential inducers for ecDNA formation include DNA repair defect (like HRD), telomere shortening, cell cycle defects, and most of these ecDNA inducers are cell-intrinsic defects.

The detailed molecular mechanism responsible for the decreased MHC class I and II antigen presentation genes’ expression, and immune evasion in ecDNA-containing tumors is not clear. The ecDNA associated oncogene could be a potential mechanism. The function of nuclear circular DNA on immune response is unknown. Cytoplasmic DNA is known to stimulate immune response through cGAS-STING pathway^28^, and in tumors with ecDNA, this pathway is not over-activated (Supplementary Fig. S7). EcDNA formation is a type of genomic DNA copy number alteration, its detections with copy number signature analysis could reveal potentially actionable biomarkers for cancer precision therapy^29–31^. Tumors with ecDNA are known to have poorer prognosis compared with tumors without ecDNA^17^. Stimulating the antigen presentation pathway could potentially revert the ecDNA-mediated immune escape.

## Materials and Methods

### Data Source

EcDNA status information was determined using AmpliconArchitect from whole genome sequencing (WGS) data as described previously ^17^. Gene expression data are available for the majority of the cancer genome atlas (TCGA) but not pan-cancer analysis of whole genomes (PCAWG) datasets. For downstream immune infiltration and gene expression analysis, we only keep TCGA samples. Tumor immune cell infiltration information for TCGA samples was downloaded from the TIMER webserver (http://timer.comp-genomics.org/), including the results calculated by TIMER, CIBERSORT, quanTIseq, xCell, and MCP-counter algorithms. Somatic mutation data detected by Mutect2 was download from UCSC xena (GDC-PANCAN.mutect2_snv.tsv). The pan-cancer gene-level RNA-Seq data of TCGA samples was downloaded from UCSC xena, including counts and normalized transcripts per million (TPM) data. Immune subtyping and tumor microenvironment (TME) information of TCGA samples are based on reports of Thorsson et al and Bagaev et al study respectively ^24,25^. The leukocyte fraction data of TCGA samples are based on the results of Thorsson et al study ^25^. In the downstream analysis, we only keep cancer types where the count of ecDNA samples was more than 20. All methods were performed in accordance with the relevant guidelines and regulations.

### Calculation of cancer immune scores

In addition to immune cell infiltration quantification, we calculated a variety of additional immune microenvironment quantitative scores. The immunophenoscore (IPS) was used to measure the immune state of the samples. IPS was based on the expression of major determinants, identified by a random forest approach, and these factors were classified into four categories: major histocompatibility complex (MHC) molecules, effector cells, suppressor cells and checkpoint markers. We used R scripts and IPS genes provided by the origin paper to calculate IPS scores ^32^. ESTIMATE (Estimation of STromal and Immune cells in MAlignant Tumor tissues using Expression data) is a tool using gene signatures to generate three scores: stromal score, immune score and estimate score, we used R package Estimate to calculate the immune score ^23^. The cytolytic activity (CYT) score was a quantitative means of assessing cytotoxic T cell infiltration and activity and was calculated as the geometric mean of expression of *GZMA* and *PRF1* genes ^33^. The tumor inflammation signature (TIS) uses 18-gene signature to measure a pre-existing but suppressed adaptive immune response within tumors. The TIS has been shown to enrich for patients who respond to the anti-PD1 agent pembrolizumab. TIS was calculated by gene set variation analysis (GSVA) using the 18-gene signature mentioned by Danaher et al^34^.

### Tumor mutational burden (TMB) and neoantigen burden

TMB was defined as the number of non-synonymous alterations per megabase (Mb) of genome examined. We used 38 Mb as the estimate of the exome size: TMB = (whole exome missense mutations) / 38. Tumor neoantigen are generated by somatic mutations, and can be recognized as foreign by immune cells, conferring immunogenicity to cancer cells. Neoantigen was predicted based on somatic mutation and human leukocyte antigen (HLA) typing data. HLA typing data for TCGA cancer was obtained from Thorsson et al study^25^. Mutect2 mutation files were first transformed into VCF format by maf2vcf tools, and we used NeoPredPipe to predict neoantigen^35^. We only evaluated single-nucleotide variants leading to a single amino acid change, and novel peptides of nine amino acids were considered. From the output results, if the IC50 of a novel peptide is less than 50nM, and the TPM expression level is greater than 1, then this peptide is labeled as neoantigen. A mutation was considered neoantigenic if there was at least a single peptide produced from the mutated base that produce a neoantigen. Neoantigen burden was calculated similarly as TMB: (Total counts of neoantigens in the exome) / 38.

### Gene set enrichment analysis (GSEA)

For each cancer type, we used Deseq2 to identify differentially expressed genes between ecDNA and non-ecDNA samples^36^. Then gene set enrichment analysis was performed by using R package “fgsea”. We downloaded gene list gmt file for the following pathways from MSigDB database, including “REACTOME_MHC_CLASS_II_ANTIGEN_PRESENTATION”, “REACTOME_CLASS_I_MHC_MEDIATED_ANTIGEN_PROCESSING_PRE SENTATION”, “GOBP_ANTIGEN_PROCESSING_AND_PRESENTATION_OF_PEPTIDE_ANTIGEN_VIA_MHC_CLASS_I”, and “GOBP_ANTIGEN_PROCESSING_AND_PRESENTATION_OF_PEPTIDE_OR_POLYSACCHARIDE_ANTIGEN_VIA_MHC_CLASS_II”. The GSEA p-values were corrected by FDR method, and was considered significant if less than 0.05. For each cancer sample, we also calculated corresponding pathway GSVA scores using R package “GSVA”^37^.

### Statistical analysis

All *P* values showed in boxplot were calculated by Wilcoxon tests using R. We used the following convention for symbols indicating statistical significance: ns: *P*>0.05, *: *P*≤0.05, **: *P*≤0.01, ***: *P*≤0.001, ****: *P*≤0.0001. Immune subtype enrichment analysis was conducted by chi-squared test. All statistical tests and visualization analyses were performed with R.

### Data Availability Statement

Only publicly available data were used in this study, and data sources and handling of these data are described in the Materials and Methods and in Supplementary Table 1-3. All codes required to reproduce the results reported in this manuscript are freely available at: https://github.com/XSLiuLab/ecDNA_immune. Analyses can be read online at: https://xsliulab.github.io/ecDNA_immune/. Further information is available from the corresponding author upon request.

## Acknowledgements

We thank ShanghaiTech University High Performance Computing Public Service Platform for computing services. We thank Raymond Shuter for editing the text. We thank multi-omics facility, molecular and cell biology core facility of ShanghaiTech University for technical help. This work was supported by Shanghai Science and Technology Commission (21ZR1442400), the National Natural Science Foundation of China (31771373), and startup funding from ShanghaiTech University.

## Contributions

TW, CW, collected the data and performed the computational analysis. XZ, GW, WN, ZT, FC participated in critical project discussion. XSL designed, supervised the study and wrote the manuscript.

