## Supplemental Figures for "Extrachromosomal DNA is associated with decreased immune cell infiltration and antigen presentation, represents a potential cancer immune evasion mechanism"

**Supplementary Table 1: List of datasets and software used in this study**

| data/software | resource |
|---|---|
| TCGA somatic mutation | https://gdc-hub.s3.us-east-1.amazonaws.com/download/GDC-PANCAN.mutect2_snv.tsv.gz |
| TCGA mRNA expression | https://toil-xena-hub.s3.us-east-1.amazonaws.com/download/tcga_RSEM_gene_tpm.gz |
| TCGA ecDNA information | Kim H, Nguyen N P, Turner K, et al. Extrachromosomal DNA is associated with oncogene amplification and poor outcome across multiple cancers[J]. Nature genetics, 2020, 52(9): 891-897. |
| ESTIMATE | R estimate package: https://bioinformatics.mdanderson.org/estimate/rpackage.html |
| Immune cell infiltration estimation | TIMER webserver: http://timer.comp-genomics.org/, including the results calculated by TIMER, CIBERSORT, quanTIseq, xCell, and MCP-counter algorithms |
| Leukocyte fraction | Thorsson V, Gibbs D L, Brown S D, et al. The immune landscape of cancer[J]. Immunity, 2018, 48(4): 812-830. e14. (https://gdc.cancer.gov/about-data/publications/panimmune) |
| Immunophenoscore (IPS) | Charoentong P, Finotello F, Angelova M, et al. Pan-cancer immunogenomic analyses reveal genotype-immunophenotype relationships and predictors of response to checkpoint blockade[J]. Cell reports, 2017, 18(1): 248-262. |
| Tumor inflammation signature (TIS) | Danaher P, Warren S, Lu R, et al. Pan-cancer adaptive immune resistance as defined by the Tumor Inflammation Signature (TIS): results from The Cancer Genome Atlas (TCGA)[J]. Journal for immunotherapy of cancer, 2018, 6(1): 1-17. Use GSVA to calculate TIS. |
| Cytolytic activity score (CYT) | Geometric mean of expression of GZMA and PRF1 genes |
| TCGA samples immune subtype | 1.Thorsson V, Gibbs D L, Brown S D, et al. The immune landscape of cancer[J]. Immunity, 2018, 48(4): 812-830. e14. (https://gdc.cancer.gov/about-data/publications/panimmune) 2.Bagaev A, Kotlov N, Nomie K, et al. Conserved pan-cancer microenvironment subtypes predict response to immunotherapy[J]. Cancer Cell, 2021, 39(6): 845-865. e7. |
| maf2vcf | https://github.com/mskcc/vcf2maf/blob/main/maf2vcf.pl |
| neoantigen prediction software: NeoPredPipe | https://github.com/MathOnco/NeoPredPipe |

### Supplementary Table 2: List of genes in gene sets used for GSVA

| pathways | genes for GSVA analysis |
|---|---|
| GO MHC-I | ABCB9,ACE,AZGP1,B2M,BCAP31,CALR,CANX,CD207,CD36,CHUK,CLEC4A,CYBA,CYBB,ERAP1,ERAP2,FCER1G,FCGR1A,FCGR1B,HFE,HLA-A,HLA-B,HLA-C,HLA-E,HLA-F,HLA-G,HLA-H,IDE,IFI30,IKBKB,IKBKG,ITGAV,ITGB5,LNPEP,MFSD6,MR1,NCF1,NCF2,NCF4,PDIA3,PSMA1,PSMA2,PSMA3,PSMA4,PSMA5,PSMA6,PSMA7,PSMA8,PSMB1,PSMB10,PSMB11,PSMB2,PSMB3,PSMB4,PSMB5,PSMB6,PSMB7,PSMB8,PSMB9,PSMC1,PSMC2,PSMC3,PSMC4,PSMC5,PSMC6,PSMD1,PSMD10,PSMD11,PSMD12,PSMD13,PSMD14,PSMD2,PSMD3,PSMD4,PSMD5,PSMD6,PSMD7,PSMD8,PSMD9,PSME1,PSME2,PSME3,PSME4,PSMF1,SAR1B,SEC13,SEC22B,SEC23A,SEC24A,SEC24B,SEC24C,SEC24D,SEC31A,SNAP23,TAP1,TAP2,TAPBP,TAPBPL,VAMP3,VAMP8 |
| GO MHC-II | ACTR10,ACTR1A,ACTR1B,AP1B1,AP1G1,AP1M1,AP1M2,AP1S1,AP1S2,AP1S3,AP2A1,AP2A2,AP2B1,AP2M1,AP2S1,ARF1,CANX,CAPZA1,CAPZA2,CAPZA3,CAPZB,CD74,CENPE,CLTA,CLTC,CTSD,CTSE,CTSF,CTSL,CTSS,CTSV,DCTN1,DCTN2,DCTN3,DCTN4,DCTN5,DCTN6,DNM2,DYNC1H1,DYNC1I1,DYNC1I2,DYNC1LI1,DYNC1LI2,DYNLL1,DYNLL2,FCER1G,FCGR2B,HLA-DMA,HLA-DMB,HLA-DOA,HLA-DOB,HLA-DPA1,HLA-DPB1,HLA-DQA1,HLA-DQA2,HLA-DQB1,HLA-DQB2,HLA-DRA,HLA-DRB1,HLA-DRB3,HLA-DRB4,HLA-DRB5,IFI30,KIF11,KIF15,KIF18A,KIF22,KIF23,KIF26A,KIF2A,KIF2B,KIF2C,KIF3A,KIF3B,KIF3C,KIF4A,KIF4B,KIF5A,KIFAP3,KLC1,KLC2,LAG3,LGMN,MARCHF1,MARCHF8,OSBPL1A,PIKFYVE,PYCARD,RAB7A,RACGAP1,RILP,SAR1B,SEC13,SEC23A,SEC24A,SEC24B,SEC24C,SEC24D,SEC31A,SH3GL2,SPTBN2,THBS1,TRAF6,TREM2 |
| REATOME MHC-I | ANAPC1,ANAPC10,ANAPC11,ANAPC13,ANAPC2,ANAPC4,ANAPC5,ANAPC7,AREL1,ARIH2,ASB1,ASB10,ASB11,ASB12,ASB13,ASB14,ASB15,ASB16,ASB17,ASB18,ASB2,ASB3,ASB4,ASB5,ASB6,ASB7,ASB8,ASB9,ATG7,B2M,BCAP31,BLMH,BTBD1,BTBD6,BTK,BTRC,CALR,CANX,CBLB,CBLL2,CCNF,CD14,CD207,CD36,CDC16,CDC20,CDC23,CDC26,CDC27,CDC34,CHUK,CTSL,CTSS,CTSV,CUL1,CUL2,CUL3,CUL5,CUL7,CYBA,CYBB,DCAF1,DET1,DTX3L,DZIP3,ELOB,ELOC,ERAP1,ERAP2,FBXL12,FBXL13,FBXL14,FBXL15,FBXL16,FBXL18,FBXL19,FBXL20,FBXL22,FBXL3,FBXL4,FBXL5,FBXL7,FBXL8,FBXO10,FBXO11,FBXO15,FBXO17,FBXO2,FBXO21,FBXO22,FBXO27,FBXO30,FBXO31,FBXO32,FBXO4,FBXO40,FBXO41,FBXO44,FBXO6,FBXO7,FBXO9,FBXW10,FBXW11,FBXW12,FBXW2,FBXW4,FBXW5,FBXW7,FBXW8,FBXW9,FCGR1A,FCGR1B,FGA,FGB,FGG,FZR1,GAN,GLMN,HACE1,HECTD1,HECTD2,HECTD3,HECW2,HERC1,HERC2,HERC3,HERC4,HERC5,HERC6,HLA-A,HLA-B,HLA-C,HLA-E,HLA-F,HLA-G,HMGB1,HSPA5,HUWE1,IKBKB,IKBKG,ITCH,ITGAV,ITGB5,KBTBD13,KBTBD6,KBTBD7,KBTBD8,KCTD6,KCTD7,KEAP1,KLHL11,KLHL13,KLHL2,KLHL20,KLHL21,KLHL22,KLHL25,KLHL3,KLHL41,KLHL42,KLHL5,KLHL9,LMO7,LNPEP,LNX1,LONRF1,LRR1,LRRC41,LRSAM1,LTN1,LY96,MEX3C,MGRN1,MIB2,MKRN1,MRC1,MRC2,MYD88,MYLIP,NCF1,NCF2,NCF4,NEDD4,NEDD4L,NPEPPS,PDIA3,PJA1,PJA2,PRKN,PSMA1,PSMA2,PSMA3,PSMA4,PSMA5,PSMA6,PSMA7,PSMA8,PSMB1,PSMB10,PSMB11,PSMB2,PSMB3,PSMB4,PSMB5,PSMB6,PSMB7,PSMB8,PSMB9,PSMC1,PSMC2,PSMC3,PSMC4,PSMC5,PSMC6,PSMD1,PSMD10,PSMD11,PSMD12,PSMD13,PSMD14,PSMD2,PSMD3,PSMD4,PSMD5,PSMD6,PSMD7,PSMD8,PSMD9,PSME1,PSME2,PSME3,PSME4,PSMF1,RBBP6,RBCK1,RBX1,RCHY1,RLIM,RNF111,RNF114,RNF115,RNF123,RNF126,RNF130,RNF138,RNF14,RNF144B,RNF182,RNF19A,RNF19B,RNF213,RNF217,RNF220,RNF25,RNF34,RNF4,RNF41,RNF6,RNF7,RPS27A,S100A1,S100A8,S100A9,SAR1B,SEC13,SEC22B,SEC23A,SEC24A,SEC24B,SEC24C,SEC24D,SEC31A,SEC61A1,SEC61A2,SEC61B,SEC61G,SEM1,SH3RF1,SIAH1,SIAH2,SKP1,SKP2,SMURF1,SMURF2,SNAP23,SOCS1,SOCS3,SPSB1,SPSB2,SPSB4,STUB1,STX4,TAP1,TAP2,TAPBP,THOP1,TIRAP,TLR1,TLR2,TLR4,TLR6,TPP2,TRAF7,TRAIP,TRIM11,TRIM21,TRIM32,TRIM36,TRIM37,TRIM39,TRIM4,TRIM41,TRIM50,TRIM63,TRIM69,TRIM71,TRIM9,TRIP12,UBA1,UBA3,UBA5,UBA52,UBA6,UBA7,UBAC1,UBB,UBC,UBE2A,UBE2B,UBE2C,UBE2D1,UBE2D2,UBE2D3,UBE2D4,UBE2E1,UBE2E2,UBE2E3,UBE2F,UBE2G1,UBE2G2,UBE2H,UBE2J1,UBE2J2,UBE2K,UBE2L3,UBE2L6,UBE2M,UBE2N,UBE2O,UBE2Q1,UBE2Q2,UBE2R2,UBE2S,UBE2U,UBE2V1,UBE2V2,UBE2W,UBE2Z,UBE3A,UBE3B,UBE3C,UBE3D,UBE4A,UBOX5,UBR1,UBR2,UBR4,UFL1,UNKL,VAMP3,VAMP8,VHL,WSB1,WWP1,ZBTB16,ZNRF1,ZNRF2 |
| REATOME MHC-II | ACTR10,ACTR1A,ACTR1B,AP1B1,AP1G1,AP1M1,AP1M2,AP1S1,AP1S2,AP1S3,AP2A1,AP2A2,AP2B1,AP2M1,AP2S1,ARF1,CANX,CAPZA1,CAPZA2,CAPZA3,CAPZB,CD74,CENPE,CLTA,CLTC,CTSA,CTSB,CTSC,CTSD,CTSE,CTSF,CTSH,CTSK,CTSL,CTSO,CTSS,CTSV,DCTN1,DCTN2,DCTN3,DCTN4,DCTN5,DCTN6,DNM1,DNM2,DNM3,DYNC1H1,DYNC1I1,DYNC1I2,DYNC1LI1,DYNC1LI2,DYNLL1,DYNLL2,HLA-DMA,HLA-DMB,HLA-DOA,HLA-DOB,HLA-DPA1,HLA-DPB1,HLA-DQA1,HLA-DQA2,HLA-DQB1,HLA-DQB2,HLA-DRA,HLA-DRB1,HLA-DRB3,HLA-DRB4,HLA-DRB5,IFI30,KIF11,KIF15,KIF18A,KIF20A,KIF22,KIF23,KIF26A,KIF2A,KIF2B,KIF2C,KIF3A,KIF3B,KIF3C,KIF4A,KIF4B,KIF5A,KIF5B,KIFAP3,KLC1,KLC2,KLC3,KLC4,LAG3,LGMN,OSBPL1A,RAB7A,RACGAP1,RILP,SAR1B,SEC13,SEC23A,SEC24A,SEC24B,SEC24C,SEC24D,SEC31A,SH3GL2,SPTBN2,TUBA1A,TUBA1B,TUBA1C,TUBA3C,TUBA3D,TUBA3E,TUBA4A,TUBA4B,TUBA8,TUBAL3,TUBB1,TUBB2A,TUBB2B,TUBB3,TUBB4A,TUBB4B,TUBB6,TUBB8,TUBB8B |

**Supplementary Table 3: Number of sample information**

| | cancer type | ecDNA positive | ecDNA negative |
|---|---|---|---|
| Number of TCGA samples with WGS data available for ecDNA detection | BLCA | 33 | 79 |
| | BRCA | 29 | 79 |
| | CESC | 15 | 51 |
| | COAD | 4 | 52 |
| | DLBC | 1 | 6 |
| | ESCA | 29 | 31 |
| | GBM | 28 | 19 |
| | HNSC | 39 | 114 |
| | KICH | 1 | 49 |
| | KIRC | 1 | 43 |
| | KIRP | 2 | 34 |
| | LAML | 0 | 40 |
| | LGG | 11 | 74 |
| | LIHC | 4 | 48 |
| | LUAD | 22 | 121 |
| | LUSC | 14 | 36 |
| | OV | 11 | 34 |
| | PRAD | 2 | 119 |
| | READ | 0 | 18 |
| | SARC | 17 | 19 |
| | SKCM | 23 | 113 |
| | STAD | 32 | 95 |
| | THCA | 0 | 136 |
| | UCEC | 20 | 123 |
| | UVM | 1 | 49 |
| Number of TCGA samples with gene expression data available | BLCA | 33 | 79 |
| | BRCA | 29 | 79 |
| | CESC | 15 | 50 |
| | COAD | 4 | 29 |
| | DLBC | 1 | 6 |
| | ESCA | 28 | 30 |
| | GBM | 18 | 11 |

| | cancer type | ecDNA positive | ecDNA negative |
|---|---|---|---|
| | HNSC | 39 | 113 |
| | KICH | 1 | 49 |
| | KIRC | 1 | 43 |
| | KIRP | 2 | 34 |
| | LAML | 0 | 31 |
| | LGG | 11 | 72 |
| | LIHC | 4 | 46 |
| | LUAD | 22 | 119 |
| | LUSC | 14 | 36 |
| | OV | 10 | 25 |
| | PRAD | 2 | 119 |
| | READ | 0 | 2 |
| | SARC | 17 | 19 |
| | SKCM | 23 | 113 |
| | STAD | 27 | 87 |
| | THCA | 0 | 133 |
| | UCEC | 1 | 8 |
| | UVM | 1 | 48 |

Figure S1

a

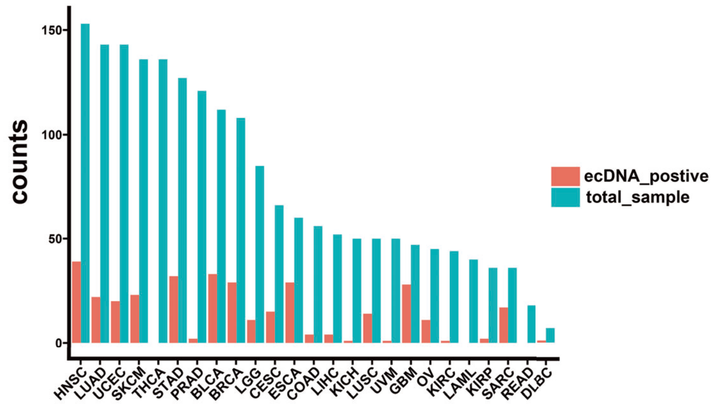


b

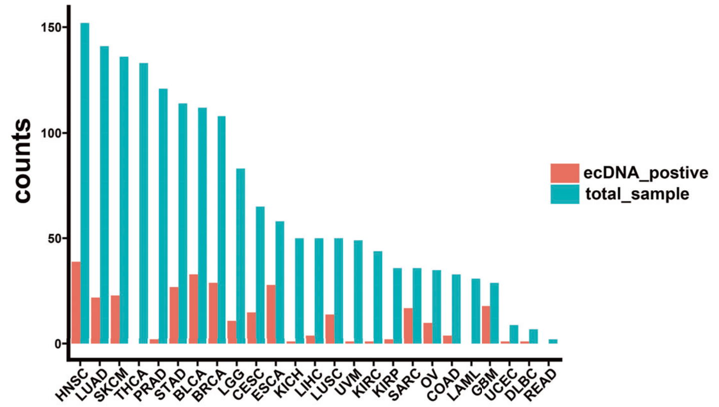


**Supplementary Fig. S1. Sample counts distribution of the cancer types with WGS and gene expression data available for ecDNA analysis.**

a. Total number of cancer samples with ecDNA status information.

b. Number of cancer samples with both ecDNA status and gene expression information. TCGA cancer type acronyms: THYM (thymoma), ESCA (esophageal carcinoma), BRCA (breast invasive carcinoma), LUAD (lung adenocarcinoma), LUSC (lung squamous cell carcinoma), KICH (kidney chromophobe), STAD (stomach adenocarcinoma), CHOL (cholangiocarcinoma), LIHC (liver hepatocellular carcinoma), PRAD (prostate adenocarcinoma), HNSC (head and neck squamous cell carcinoma), KIRP (kidney renal papillary cell carcinoma), SARC (sarcoma), UCEC (uterine corpus endometrial carcinoma), BLCA (bladder urothelial carcinoma), PAAD (pancreatic adenocarcinoma), CESC (cervical squamous cell carcinoma and endocervical adenocarcinoma), GBM (glioblastoma multiforme), KIRC (kidney renal clear cell carcinoma), SKCM (skin cutaneous melanoma), PCPG (pheochromocytoma and paraganglioma), THCA (thyroid carcinoma), LGG (Brain Lower Grade Glioma), UVM (Uveal Melanoma), OV (Ovarian serous cystadenocarcinoma), COAD(Colon adenocarcinoma), LAML (Acute Myeloid Leukemia), DLBC (Lymphoid Neoplasm Diffuse Large B-cell Lymphoma), READ (Rectum adenocarcinoma).

Figure S2

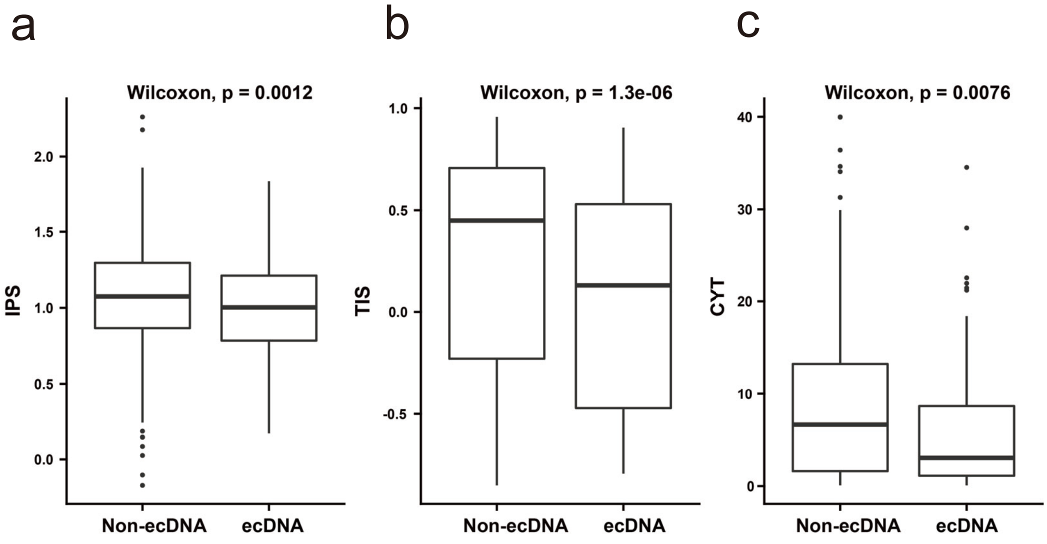


**Supplementary Fig. S2 Comparisons of immune infiltration scores calculated by different methods between ecDNA and non-ecDNA samples.**

a. Immunophenoscore (IPS) was used to measure the immune state of the samples, and was calculated according to Charoentong et al 2017 study.

b. Tumor inflammation signature (TIS) score was calculated by GSVA using the gene signature described in Danaher et al 2018 study.

c. Cytolytic activity score (CYT) was calculated as the geometric mean of expression of GZMA and PRF1 genes. Wilcoxon test p values are shown.

Figure S3

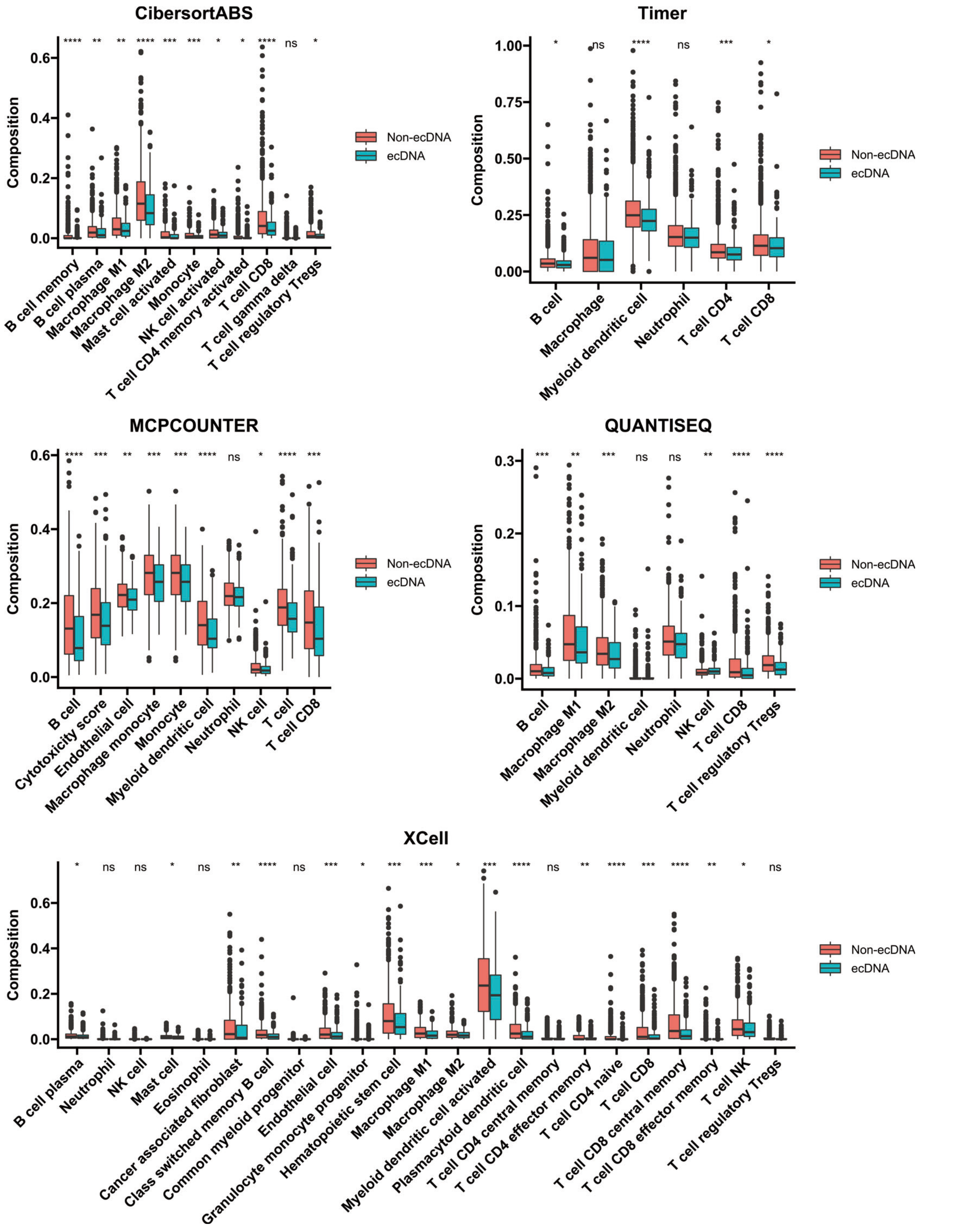


**Supplementary Fig. S3 Comparisons of immune cell compositions quantified by five different methods between ecDNA and non-ecDNA samples.**

Wilcoxon test p values are shown. ns: $p>0.05$, *: $p\leq0.05$, **: $p\leq0.01$, ***: $p \leq0.001$, ****: $p\leq 0.0001$.

Figure S4

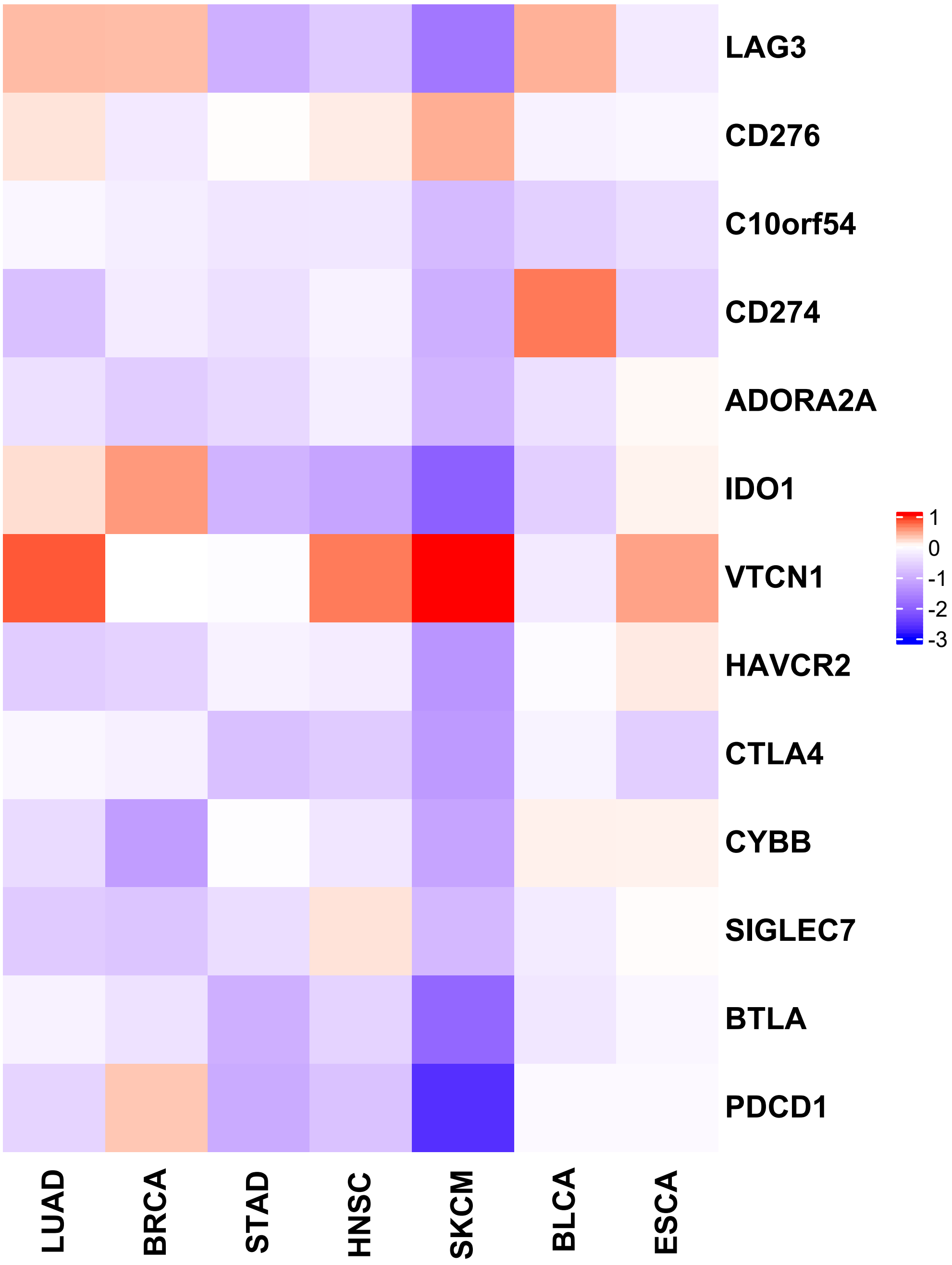


**Supplementary Fig. S4 Pan-cancer comparisons of the expression of inhibitory immune checkpoint genes between tumors with and without ecDNA.**
Heatmap color indicates the difference between the median expression for specific gene in specific cancer type between ecDNA and non-ecDNA samples, i.e. median expression of specific gene (row) in samples with ecDNA minus median expression of this gene in samples without ecDNA in a cancer type (column). TCGA cancer type acronyms: ESCA (esophageal carcinoma), BRCA (breast invasive carcinoma), LUAD (lung adenocarcinoma), STAD (stomach adenocarcinoma), HNSC (head and neck squamous cell carcinoma), BLCA (bladder urothelial carcinoma), SKCM (skin cutaneous melanoma).

Figure S5

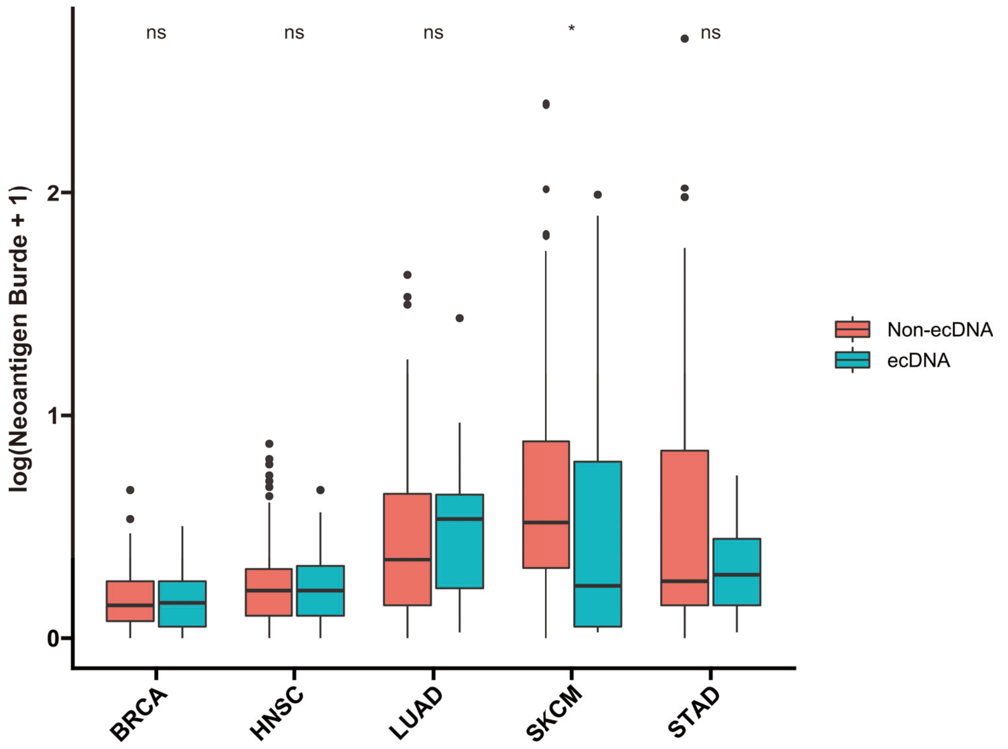


**Supplementary Fig. S5 Pan-cancer comparisons of tumor neoantigen burden in tumors with and without ecDNA.**
Wilcoxon test p values are shown. ns: $p>0.05$, *: $p\leq0.05$. TCGA cancer type acronyms: BRCA (breast invasive carcinoma), LUAD (lung adenocarcinoma), STAD (stomach adenocarcinoma), HNSC (head and neck squamous cell carcinoma), SKCM (skin cutaneous melanoma).

Figure S6

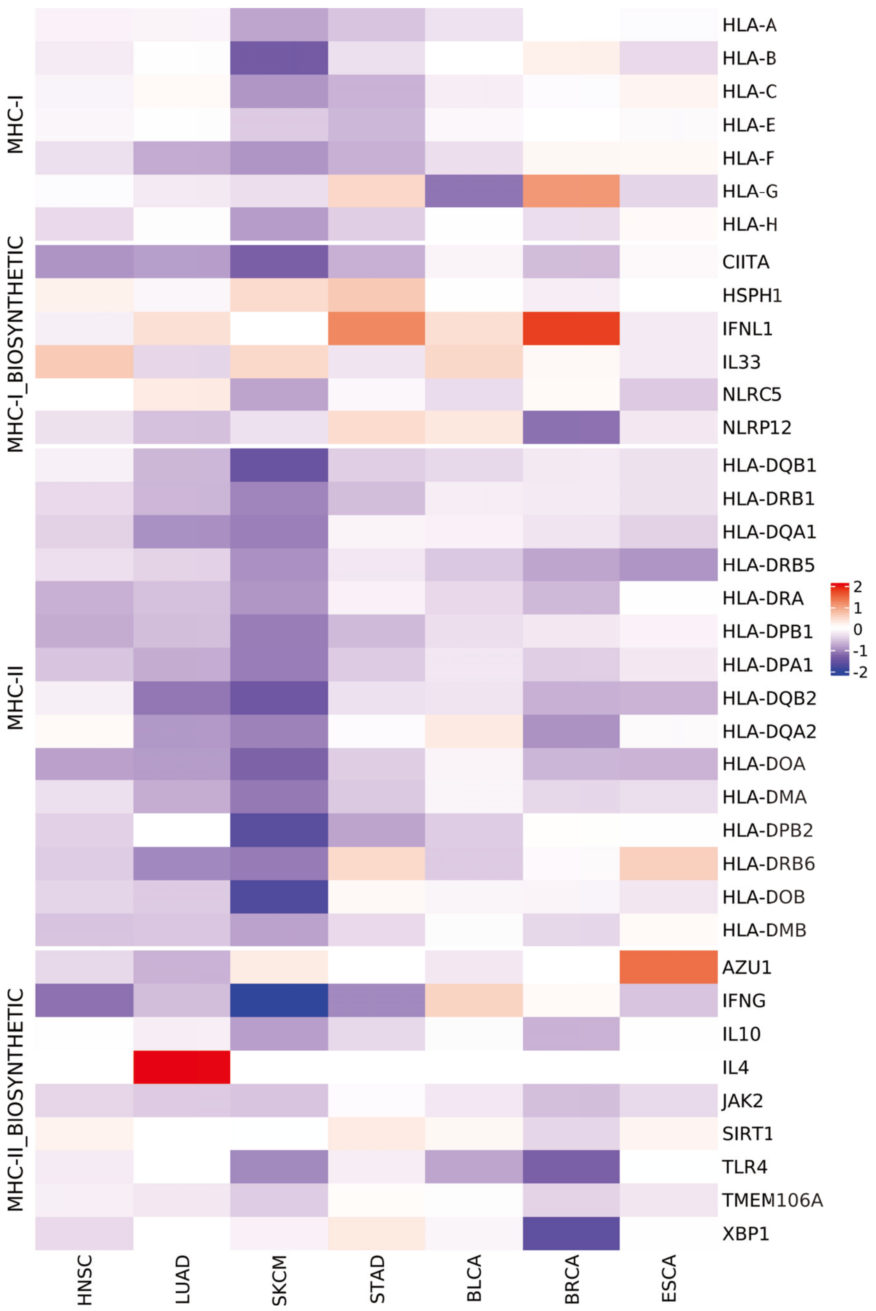


**Supplementary Fig. S6 Pan-cancer comparisons of the expression of MHC class I and class II antigen presentation genes in tumor with and without ecDNA.**
Heatmap color indicates the difference between the median expression for specific gene in specific cancer type between ecDNA and non-ecDNA samples. TCGA cancer type acronyms: ESCA (esophageal carcinoma), BRCA (breast invasive carcinoma), LUAD (lung adenocarcinoma), STAD (stomach adenocarcinoma), HNSC (head and neck squamous cell carcinoma), BLCA (bladder urothelial carcinoma), SKCM (skin cutaneous melanoma).

Figure S7

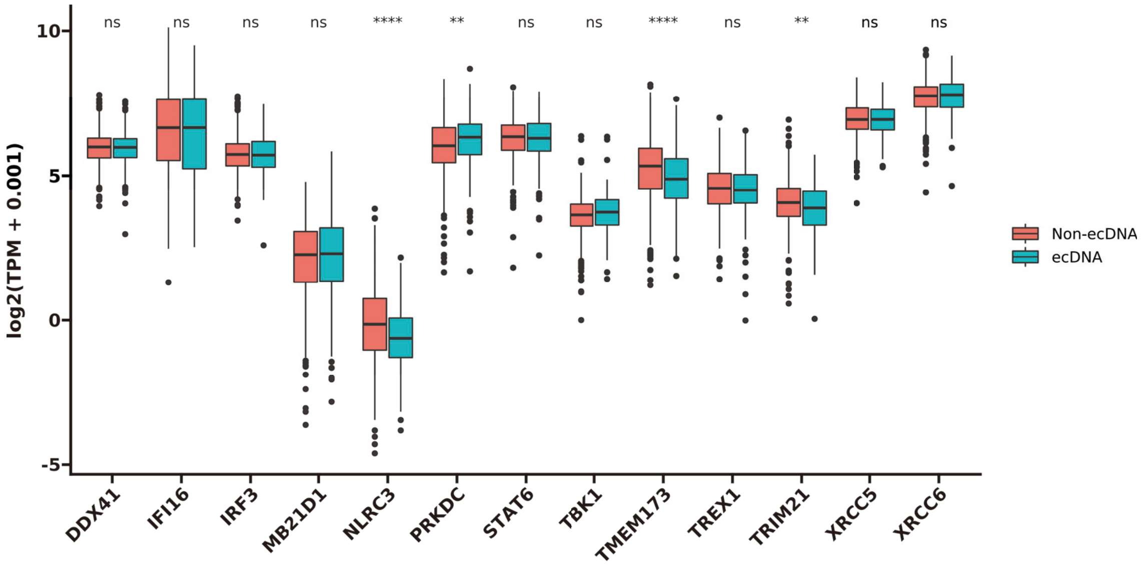


**Supplementary Fig. S7 Comparisons of the expression of cGAS-STING pathway genes between tumors with and without ecDNA.**

Wilcoxon test p values are shown. ns: $p>0.05$, *: $p\leq0.05$, **: $p\leq0.01$, ***: $p\leq0.001$, ****: $p\leq0.0001$.
